# Donor-Specific Plasma Factors Contribute to Aberrant Staining Artifacts with a Commercial Lyophilized Mass Cytometry Antibody Panel

**DOI:** 10.1101/2020.11.30.405027

**Authors:** Daniel Geanon, Brian Lee, Adeeb H. Rahman

## Abstract

High-dimensional mass cytometry (CyTOF) phenotyping allows for the routine measurement of over 40 parameters and is increasingly being utilized across a wide range of studies. However, CyTOF-specific panel design and optimization represent challenges to wider adoption and standardization of immune profiling with CyTOF. To address this, Fluidigm recently commercialized its MaxPar Direct Immune Profiling Assay (MDIPA), which comprises a lyophilized 30-marker antibody panel that is able to identify all major circulating immune cell subsets and offers a streamlined solution for standardized human immune monitoring. However, in the course of applying the MDIPA to characterize large numbers of whole blood samples, we observed several instances of unusual aberrant staining patterns, most notably CD19 expression on non-B cells, which can potentially confound data analysis and lead to erroneous interpretation of results when using this assay. Here, we report that this complex phenomenon is mediated by donor-specific plasma factors that mediate non-specific interactions between specific antibodies in the MDIPA panel. Our findings additionally suggest specific strategies that can be used to mitigate the issue, including the use of PBMCs or lysed/washed whole blood to remove endogenous plasma prior to staining, or blocking specific antibodies in the MDIPA panel.

## Introduction

Mass cytometry (CyTOF) allows for the detection of over 40 parameters with minimal signal spillover between detection channels, and is now a heavily utilized technology for high dimensional single cell characterization, yielding incredible insights into the complex heterogeneity of the human immune system. As with all antibody-targeted discovery platforms, the insights gained from mass cytometry are largely driven by the antibody panels used. The design, validation, optimization and formulation of large antibody panels represents a significant source of technical variability across mass cytometry studies and presents a challenge to wider adoption and standardization of the technology. The recent release of the Maxpar Direct Immune Profiling Assay (MDIPA) represents a significant step forward in making CyTOF more accessible to the greater immunology community [1]. The MDIPA contains a standardized lyophilized panel of 30 antibodies that are sufficient to identify all major circulating immune cell subsets, and allows for supplementation and customization with additional antibodies. The assay is extremely simple to use, requiring the addition of whole blood or PBMCs directly to the tube containing the lyophilized antibody panel. The MDIPA-stained cells can also be fixed, frozen and shipped to remote sites for data acquisition, facilitating standardized CyTOF immune profiling even in the setting of multicenter studies where local mass cytometry equipment is not available. Based on these advantages, we and others have begun utilizing the MDIPA across several large research studies. However, having applied this assay to whole blood samples from several hundred patients, we have observed aberrant antibody staining patterns in approximately 20% of patient samples. Here, we report this to be an MDIPA-specific issue resulting from interactions of specific MDIPA antibodies with donor-specific serum factors in whole blood samples.

## Methods

### Subjects

The large cohort of blood samples analyzed as part of the study were collected as part of the Mt. Sinai COVID-19 biobank [2] from hospitalized patients under an institutionally approved IRB protocol. However, to rule out a potential contribution for disease status to the artifact reported here, we focused these experiments on samples collected from two consented healthy control donors (designated as Donors A and B in the figures).

### Antibodies

The monoclonal antibodies utilized in this study are listed in Table 1.

**Table 1.** Antibodies.

| Target | Clone | Conjugate | Source |
| --- | --- | --- | --- |
| CD45 | HI30 | 89 Y | Fluidigm MDIPA |
| CCR6 | G034E3 | 141 Pr | Fluidigm MDIPA |
| CD123 | 6H6 | 143 Nd | Fluidigm MDIPA |
| CD19 | HIB19 | 144 Nd | Fluidigm MDIPA |
| CD4 | RPA-T4 | 145 Nd | Fluidigm MDIPA |
| CD8a | RPA-T8 | 146 Nd | Fluidigm MDIPA |
| CD11c | Bu15 | 147 Sm | Fluidigm MDIPA |
| CD16 | 3G8 | 148 Nd | Fluidigm MDIPA |
| CD45RO | UCHL1 | 149 Sm | Fluidigm MDIPA |
| CD45RA | HI100 | 150 Nd | Fluidigm MDIPA |
| CD161 | HP-3G10 | 151 Eu | Fluidigm MDIPA |
| CCR4 | L291H4 | 152 Sm | Fluidigm MDIPA |
| CD25 | BC96 | 153 Eu | Fluidigm MDIPA |
| CD27 | O323 | 154 Sm | Fluidigm MDIPA |
| CD57 | HCD57 | 155 Gd | Fluidigm MDIPA |
| CXCR3 | G025H7 | 156 Gd | Fluidigm MDIPA |
| CXCR5 | J252D4 | 158 Gd | Fluidigm MDIPA |
| CD28 | CD28.2 | 160 Gd | Fluidigm MDIPA |
| CD38 | HB-7 | 161 Dy | Fluidigm MDIPA |
| CD56 | NCAM16.2 | 163 Dy | Fluidigm MDIPA |
| TCR$\gamma\delta$ | B1 | 164 Dy | Fluidigm MDIPA |
| CD294 | BM16 | 166 Er | Fluidigm MDIPA |
| CCR7 | G043H7 | 167 Er | Fluidigm MDIPA |
| CD14 | 63D3 | 168 Er | Fluidigm MDIPA |
| CD3 | UCHT1 | 170 Er | Fluidigm MDIPA |
| CD20 | 2H7 | 171 Yb | Fluidigm MDIPA |
| CD66b | G10F5 | 172 Yb | Fluidigm MDIPA |
| HLADR | LN3 | 173 Yb | Fluidigm MDIPA |
| IgD | IA6-2 | 174 Yb | Fluidigm MDIPA |
| CD127 | A019D5 | 176 Yb | Fluidigm MDIPA |
| FITC | Polyclonal | 159 Tb | Southern Biotech; In-house conjugate (X8) |
| PE | PE001 | 165 Ho | Biolegend; In-house conjugate (X8) |
| APC | APC003 | 169 Tm | Biolegend; In-house conjugate (X8) |
| CD19 | REA675 | 142 Nd | Milteyi; In-house conjugate (X8) |
| CD19 | HIB19 | PE | Biolegend |
| CD45RA | HI100 | APC | Biolegend |
| CCR6 | G034E3 | FITC | Biolegend |

### Sample Processing

In experiments evaluating artifactual CD19-144Nd staining on healthy donor whole blood samples, fresh whole blood was first collected in Cell Preparation tubes (CPT) (BD Biosciences, Franklin Lakes, NJ, USA) and aliquoted directly into the Fluidigm MDIPA tube (Fluidigm, San Francisco, CA, USA). Per manufacturer’s protocols, 270uL of whole blood was added into the MDIPA tube and allowed to incubate for 30 minutes at room temperature (heparin blocking was not performed). After staining, 420uL of Prot1 Stabilizer (SmartTube Inc. San Carlos, CA, USA) was added directly to the sample and allowed to fix for 10 minutes at room temperature. Immediately following, the samples were transferred to cryovials and moved to −80°C for storage. For downstream processing, samples were thawed using the SmartTube Prot 1 Thaw/Erythrocyte Lysis protocol according to the manufacturer’s instructions. After erythrocyte lysis, immune cells were barcoded together utilizing the CyTOF Cell-ID 20-Plex Palladium Barcoding Kit (Fluidigm) following manufacturer’s instructions. Barcoded samples were washed in PBS + 0.2% BSA, pooled, and simultaneously fixed with 2.4% PFA in PBS with 0.08% saponin and 125nM Iridium intercalator for 30 minutes at room temperature, after which samples were washed and stored in PBS + 0.2% BSA until acquisition. This SmartTube-based protocol represents a slight modification of the vendor specified protocol, so to ensure that this protocol change was not contributing to the aberrant staining patterns, a subset of samples were also analyzed in parallel using the specific protocol recommended by the vendor for use with the MDIPA kit. In these cases, fresh whole blood was first collected in sodium heparin tubes, and 270uL of whole blood was directly aliquoted into the Fluidigm MDIPA tubes with heparin supplementation. After a 30 minute incubation at room temperature, the samples were fixed/erythrocyte lysed with Invitrogen Cal-Lyse (Thermo Fisher Scientific, Waltham, MA, USA) per the Fluidigm instructions. Samples were subsequently washed in PBS + 0.2% BSA and simultaneously fixed with 2.4% PFA in PBS with 0.08% saponin and 125nM Iridium intercalator for 30 minutes at room temperature, after which samples were washed and stored in PBS + 0.2% BSA until acquisition. In this workflow, whole blood staining, fixation, RBC lysis, and iridium intercalation all took place in one day.

In experiments evaluating artifactual staining between fresh whole blood and isolated peripheral blood mononuclear cells (PBMCs), whole blood was collected from a healthy donor with known aberrant MDIPA staining and an aliquot was processed utilizing the SmartTube fixation workflow as described above. PBMCs were isolated from the remaining blood in the CPT tubes as per the manufacturer’s protocol, washed, resuspended in FBS containing 10% DMSO and cryopreserved in liquid nitrogen. The cryopreserved PBMCs were thawed at 37°C and immediately transferred to warmed RPMI + 10% FBS media (Thermo Fisher Scientific, Waltham, MA, USA). Cells were washed once in RPMI + 10% FBS, then washed once in PBS + 0.2%BSA prior to MDIPA staining. For MDIPA staining, 3 million live cells were resuspended in 270uL PBS + 0.2% BSA and added to the MDIPA lyospheres. Samples were allowed to incubate for 30 minutes at room temperature (Fc receptor blocking & Rhodium-103 viability intercalation were performed simultaneously). Following surface staining, samples were washed, palladium mass-tagged, pooled, fixed, Ir-intercalated, and stored as described above.

In experiments evaluating the impact of donor-specific plasma on artifactual staining, whole blood was collected in sodium heparin vacutainer tubes from two healthy donors, one previously confirmed to exhibit aberrant staining patterns and one in which no aberrant staining was observed. One milliliter aliquots of blood were removed from each tube and processed using ammonium chloride lysis buffer (Stemcell) to lysis red blood cells, after which the remaining leukocytes were washed twice to remove residual plasma. In parallel, the remaining blood in the tubes was centrifuged to isolate cell-free plasma. The leukocytes from each donor were divided into three aliquots and resuspended in 333uL of PBS, autologous donor plasma, or reciprocal heterologous donor plasma. 270uL aliquots were then removed and added to separate MDIPA tubes from the same lot. Samples were allowed to incubate for 30 minutes at room temperature. Following surface staining, samples were washed, palladium mass-tagged, pooled, fixed, Ir-intercalated, and stored as described above.

In experiments utilizing fluorophore conjugated antibodies to mitigate artifactual antibody staining,270uL of whole blood was added to an MDIPA tube, and then immediately divided into four 50uL aliquots in tubes containing 1 test (per manufacturer’s guidelines) of fluorophore-conjugated CD19-PE, CD45RA-APC, CCR6-FITC or no antibody control. The samples were then incubated for 30 minutes at room temperature and then lysed with Cal-lyse per the manufacturer’s instructions. The samples were then washed, palladium mass-tagged, pooled, fixed, Ir-intercalated, and stored as described above.

### Data Acquisition and Processing

Prior to sample acquisition, samples were washed in Cell Acquisition Solution (CAS, Fluidigm), resuspended in CAS at a concentration of 1*10^6 cells/mL, supplemented with 10% Four-Element EQ Calibration Beads (Fluidigm), and acquired on the Helios Mass Cytometer at an event rate < 400 events/second with a modified wide bore injector [3]. Prior to analysis, routine data normalization (Fluidigm software) and sample demultiplexing [4] were undertaken. The debarcoded files were then uploaded to Cytobank for subsequent clean-up. Specifically, EQ beads (140Ce+) and bead-cell doublets (140Ce+Ir193+) were excluded from the data, as well as Gaussian doublets (Residual high / Offset high) and cross-sample multiplets identified by the Zunder et al. demultiplexing software (barcode_separation_dist low / mahalanobis_distance high). Immune cells were identified as CD45+Ir+ and gated for downstream analysis.

### Data Analysis

Major immune populations were defined by manual gating, and aberrant staining patterns were visualized as biaxial plots. In some cases, dimensionality reduction was performed on gated CD66b-non-granulocytes using viSNE analyses in Cytobank. CD19 and CCR6 were excluded as clustering parameters, but were visualized on the resulting viSNE maps to define the distribution of aberrant staining across cell types. The median intensity of CD19 was calculated on CD45RA+CD4+ T cells as a simple metric of aberrant staining across samples.

## Results

### Aberrant CD19 expression patterns observed with MDIPA-stained whole blood samples

The human CD19 antigen is type I transmembrane glycoprotein belonging to the immunoglobulin superfamily [5]. Outside of the neoplastic context, expression of CD19 is restricted to B cells and follicular dendritic cells, and among circulating immune cells CD19 expression is therefore generally regarded as a specific and canonical marker for B cells. However, when applying the MDIPA for large-scale immune monitoring efforts of whole blood samples from COVID-19 patients [6] we observed several instances of patient samples and healthy control samples where CD19 appeared to be aberrantly expressed on non-B cells, such as CD3-expressing T cells (Figure 1A). In most cases, the cells exhibiting this aberrant CD19 expression also showed aberrant co-expression of CCR6, a chemokine receptor that is also typically highly expressed on B cells (Figure 1B). These staining issues were not attributable to any known isotopic impurity or oxide related cross-talk, and the cells did not represent B cell aggregates or cell-cell-multiplets, as evidenced by lack of co-expression of other canonical B cell markers, such as CD20 (Figure 1C). The aberrant staining patterns appeared to be restricted to CD19 and CCR6, with all other markers in the MDIPA panel showing their expected cell-specific distributions.

**Figure 1.**
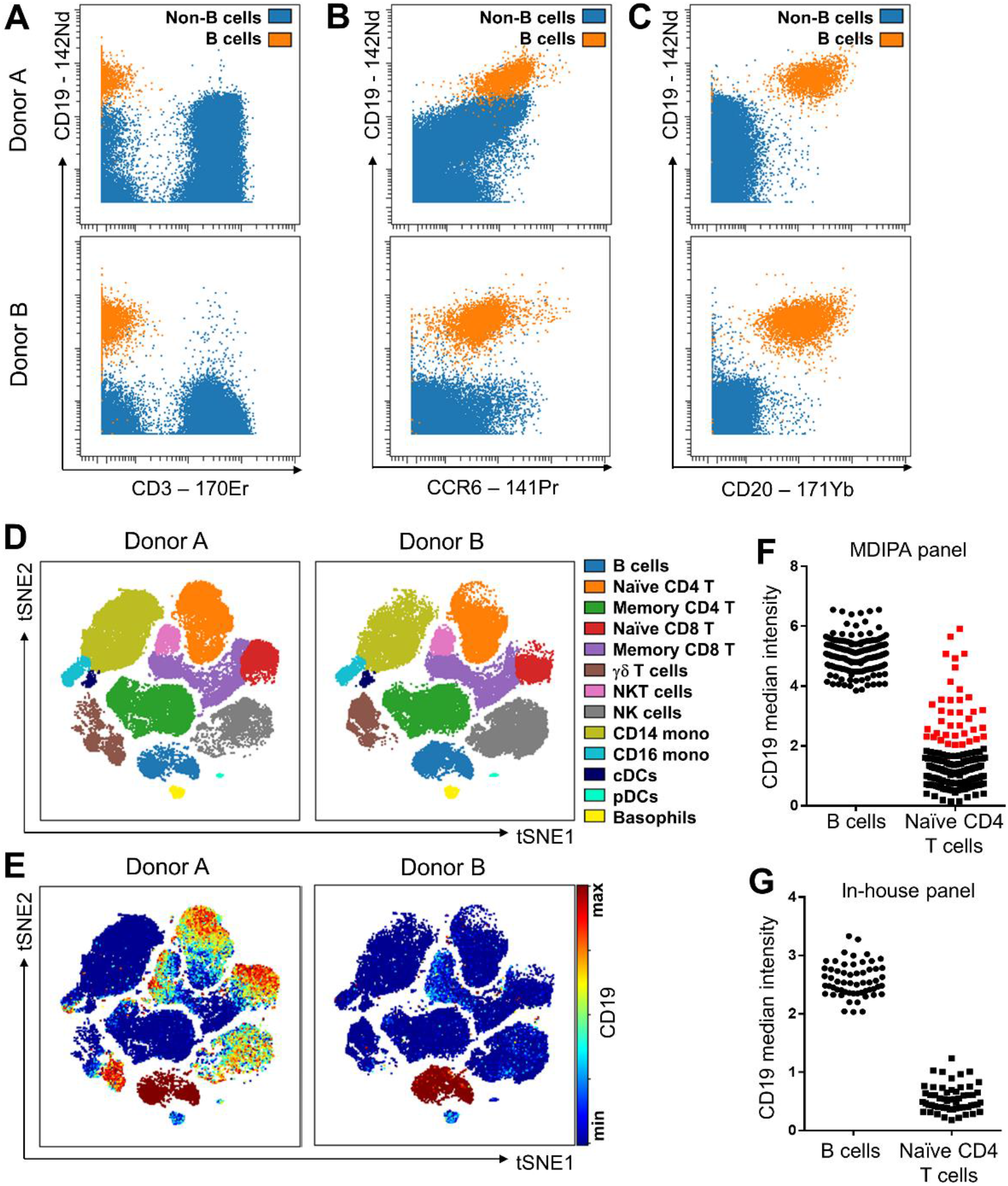
A donor-dependent CD19 staining artifact in MDIPA-stained whole blood samples. (A) Representative data from two donors illustrating the presence of aberrant CD19-144Nd staining on non-B cells in an affected donor (Donor A) in comparison to a non-affected donor (Donor B). The populations showing aberrant CD19 staining also show co-expression of CCR6 (B), however they do not show concurrent CD20 staining, suggesting that they do not represent B cell-aggregates (C). (D-E) tSNE visualization of CD19 staining across major immune subsets illustrates apparent CD19 expression across several non-B cell populations in the affected donor, most notably naive T cell populations. (F) Arcsinh-transformed median signal intensity of CD19 on B cells and naive CD4 T cells from 182 MDIPA-stained whole blood samples illustrates the donor dependency and prevalence of the staining artifact. Red dots highlight samples showing relative high levels of CD19 staining on Naive CD4 T cells (asinh-transformed median CD19 > 2), which comprise 20% of the total samples. (G) Arcsinh-transformed median signal intensity of CD19 on B cells and naive CD4 T cells from blood samples from a second cohort of 80 patients stained with an analogous workflow using an in-house conjugated lyophilized antibody panel shows no evidence of CD19 staining on T cells.

In affected individuals, unusual CD19 expression was observed across several distinct immune cell types, most obviously naive CD4 and CD8 T cells, NK cells and gamma-delta T cells (Figure 1D-E). Evaluating median CD19 expression on naive CD4 T cells therefore offered a relatively straightforward metric to determine whether or not a sample was affected by this phenomenon. When applying this metric across 182 MDIPA-stained whole blood samples, we observed varying degrees of aberrant staining across samples, with approximately 20% of samples showing high levels of staining on naive CD4 T cells (as defined by an arcsinh-transformed median intensity >2; Figure 1F). Samples with aberrant CD19 staining typically also showed higher median intensity of CD19 on B cells. These samples were processed with several different MDIPA kit lots, and while magnitude of aberrant staining seems to be somewhat affected by the specific MDIPA lots, it was nevertheless seen across most lots. Importantly, however, we did not see any evidence of similar aberrant CD19 staining when evaluating patient samples stained with an in house-antibody panel (Figure 1G), suggesting that the phenomenon was specifically related to the MDIPA panel.

### Aberrant MDIPA staining is caused by donor-specific plasma factors present in whole blood

Our results suggested that the aberrant MDIPA CD19 staining artifact was specific to blood from certain individuals. To further test this, we performed repeated blood draws from one affected individual spaced several weeks apart and independently stained blood collected at each time point with the MDIPA tubes. The MDIPA tubes used at each time point spanned different lots, and while we did observe some lot-dependency in the signal intensity of the aberrant staining, it was nevertheless consistently present across all three blood draws (Figure 2A), further confirming the donor-dependent nature of the phenomenon. Remarkably however, when PBMCs were isolated from one of the blood draws and stained with the same MDIPA panel used to stain the whole blood, we saw no evidence of the aberrant CD19 staining (Figure 2B), suggesting that this phenomenon was specific to whole blood. We therefore hypothesized that staining artifacts were related to specific factors present in the plasma of some individuals but not others. To explicitly test this hypothesis, we identified two individuals who knew to be affected (Donor A) or unaffected (Donor B) by the phenomenon based on prior whole blood staining). We collected blood from both individuals, isolated plasma and then depleted red blood cells and washed the cells to yield whole blood leukocytes that were free from plasma contamination. Consistent with our PBMC results, staining these washed whole blood leukocytes with the MDIPA panel in PBS did not show any evidence of the aberrant CD19 staining artifact, nor did staining the cells in the presence of plasma from the unaffected Donor B (Figure 2C). However, staining Donor A cells with the MDIPA in the presence of autologous Donor A plasma completely reproduced the aberrant CD19 staining phenomenon. Moreover, staining Donor B cells in the presence of heterologous Donor A plasma also resulted in the same aberrant CD19 phenomenon. These experiments definitively demonstrate that the donor dependent CD19 staining artifact is not due to intrinsic features of the leukocytes of specific individuals, but rather due to plasma factors present in the whole blood of those individuals.

**Figure 2.**
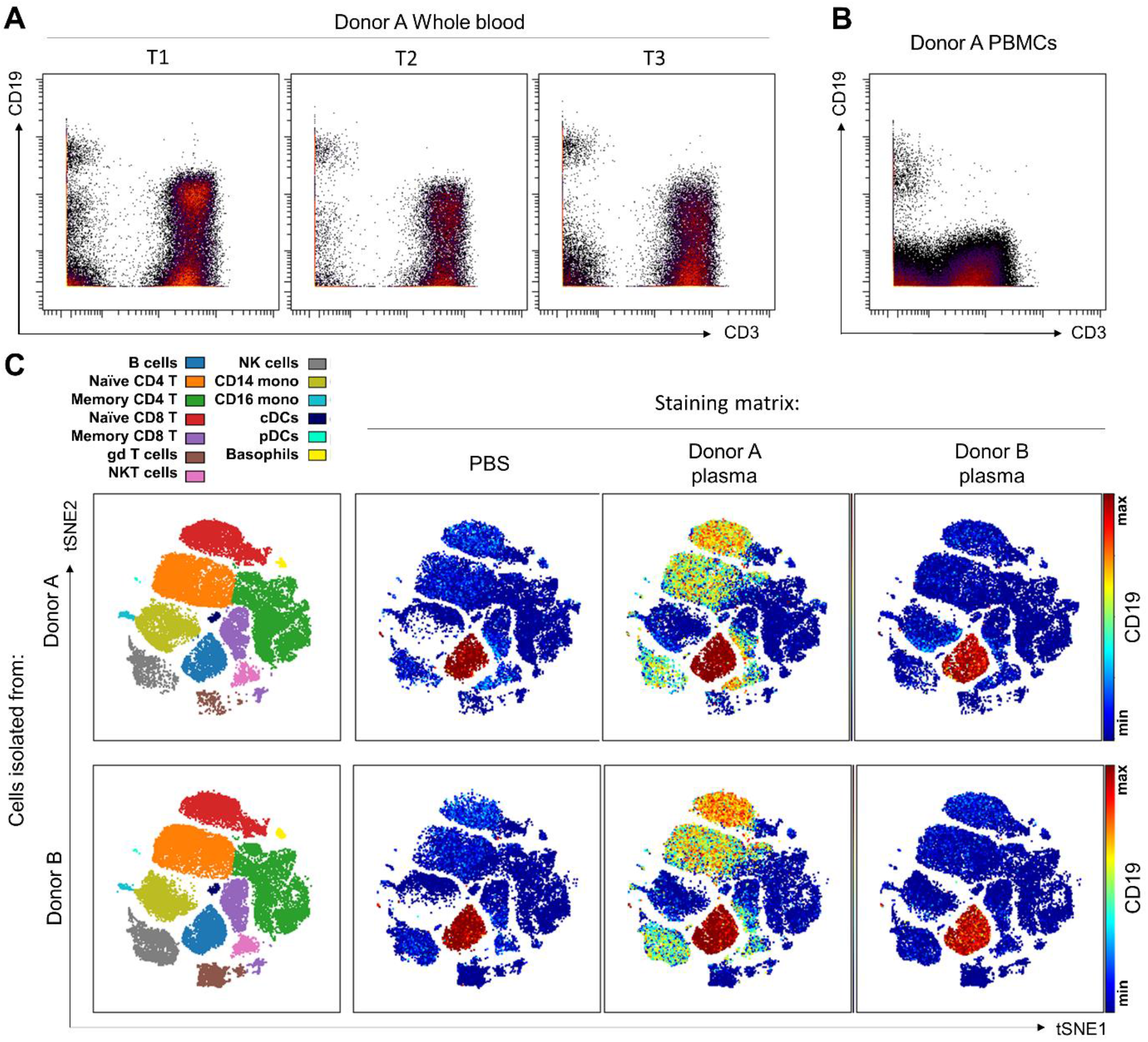
The aberrant CD19 staining artifact is specific to whole blood and mediated by donor-specific plasma factors. (A) Whole blood was collected sequentially from an affected donor on three separate occasions spaced several weeks apart and stained with independent MDIPA tubes from different lots. Biaxial plots illustrate the consistent presence of aberrant CD19 staining on non-B cells at each draw. (B) PBMCs were isolated from the same tube of blood used for the first time point whole blood stain, cryopreserved and then stained with the MDIPA panel. Biaxial plots show that aberrant CD19 is not present in the PBMCs. (C) Whole blood samples were collected from the same affected donor (Donor A) and a second donor previously confirmed to be unaffected by the staining artifact (Donor B). Plasma was collected and the cells were then RBC-lysed and washed to remove endogenous plasma. The cells from each donor were then resuspended to their original starting volume in an PBS, autologous plasma, or reciprocal heterologous plasma and stained with the MDIPA panel. Visualization of CD19 across major immune subsets highlights that no aberrant CD19 was observed in cells from either donor when stained in PBS or Donor B plasma. However, aberrant staining was restored in Donor A when stained in the presence of autologous plasma, and that this plasma could fully transfer the artifact to cells from the previously unaffected Donor B.

### The CD19 staining artifact is unrelated to the CD19 epitope or antibody paratope, but is instead mediated by interactions with CD45RA

The CD19-144Nd antibody clone present in the MDIPA panel is HIB19, which is a widely used clone that we and others have also employed across many different studies with no observation of this staining artifact. We therefore considered it unlikely that the aberrant staining pattern was due to the clone itself. To specifically evaluate this, we stained blood from an affected individual with a CD19 HIB19 antibody conjugated to PE, and then stained with the MDIPA panel. By using the same clone, we reasoned that the PE-conjugated CD19 antibody would directly compete with the CD19 antibody in the MDIPA panel for the corresponding CD19 epitope. Detecting the bound CD19-PE antibody with an anti-PE antibody showed the expected staining pattern of CD19 specific only to B cells (Figure 3A). Consistently, we observed a corresponding decrease in the MDIPA CD19-144Nd staining intensity on B cells, as would be expected from competition with the two antibodies (Figure 3B and C). Notably, however, the addition of the CD19-PE antibody had no impact on the aberrant MDIPA CD19 staining on non-B cells, indicating that this phenomenon is unrelated to the CD19 epitope or the HIB19 antibody paratope. Analogous experiments using a fluorophore-conjugated CCR6 antibody similarly confirmed that aberrant CCR6 staining was also unrelated to the CCR6 epitope of antibody paratope (data not shown).

**Figure 3.**
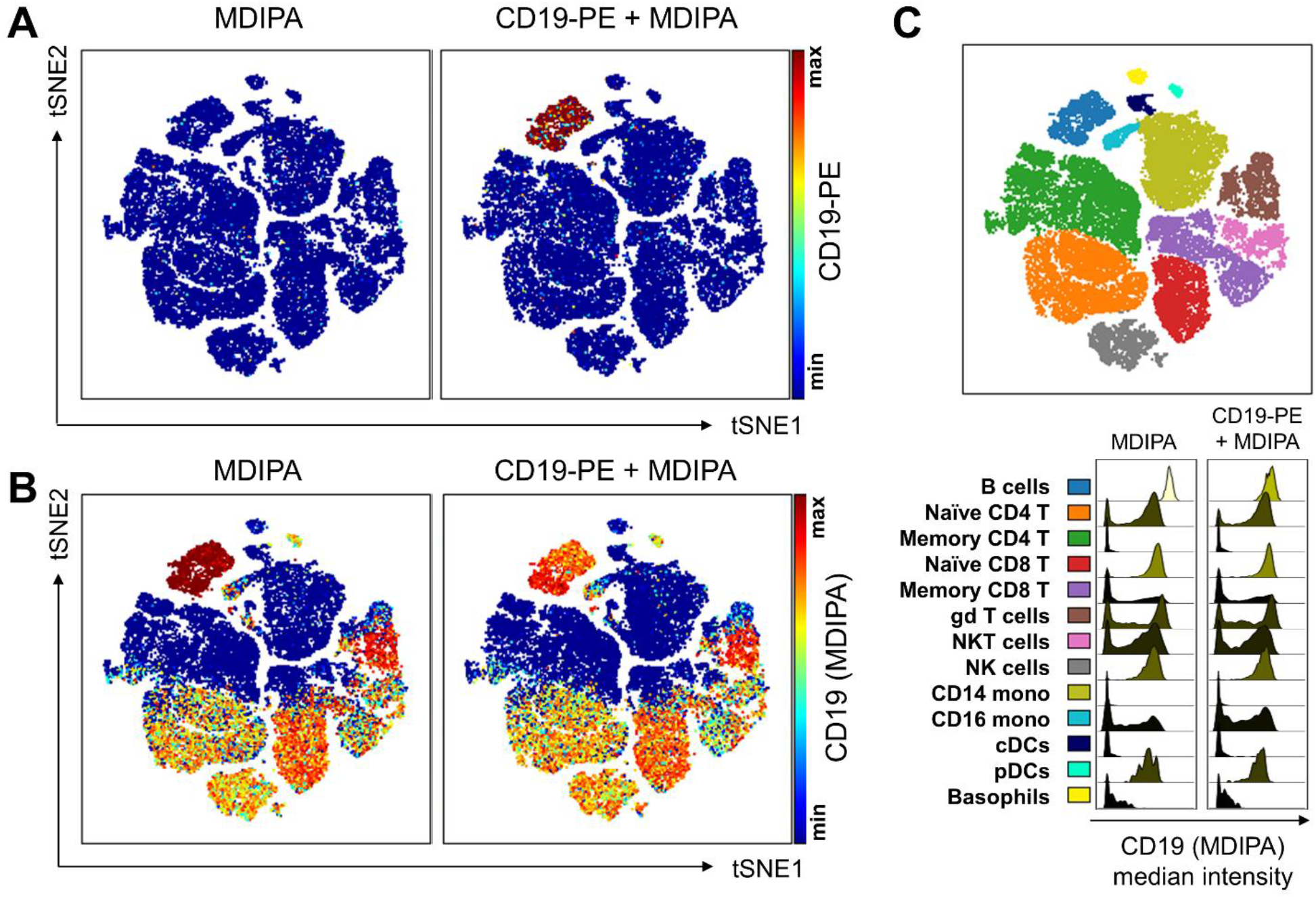
The MDIPA CD19 staining artifact is unrelated to the CD19 antibody clone, paratope or epitope. Whole blood sample from the same affected donor shown in Figures 1–3 was co-stained with the MDIPA panel and PE-conjugated CD19 clone HIB19 (the same clone contained the MDIPA panel). (A) Detection of the bound CD19-PE antibody using an anti-PE antibody shows that staining is restricted to B cells. (B-C) Addition of the PE-conjugated CD19 antibody reduces the MDIPA CD19 staining on B cells, suggesting competition for the shared CD19 epitope, but has no effect on the aberrant staining on non-B cells, suggesting that this latter staining is not mediated by an interaction between the CD19 antibody paratope and its corresponding epitope.

In performing these experiments, however, we noticed that the aberrant non-epitope associated CD19 staining appears to be highly correlated with staining of CD45RA in the MDIPA panel (Figure 4A-B). We therefore performed a similar experiment where we stained whole blood from an affected donor with a APC-conjugated CD45RA antibody matching the CD45RA antibody clone found in the MDIPA panel. This APC-conjugated antibody effectively competed and reduced staining of the CD45RA-150Nd antibody in the MDIPA panel, but detection of the same CD45RA staining pattern was faithfully reproduced by detecting the bound APC-CD45RA antibody with an anti-APC antibody (Figure 4C). Remarkably however, blocking the CD45RA antibody in the MDIPA panel also effectively blocked the aberrant CD19 staining and the aberrant CCR6 staining (Figure 4D-F). Blocking CD45RA also reduced CD19 staining intensity on B cells, highlighting that some of the apparent CD19 staining on B cells was in fact likely due to the aberrant staining of CD45RA on these cells. These findings suggest that the aberrant staining pattern is mediated by an interaction of the CD45RA, CD19 and CCR6 antibodies in the MDIPA panel with the CD45RA epitope, and that blocking this epitope consequently reduces the aberrant staining.

**Figure 4.**
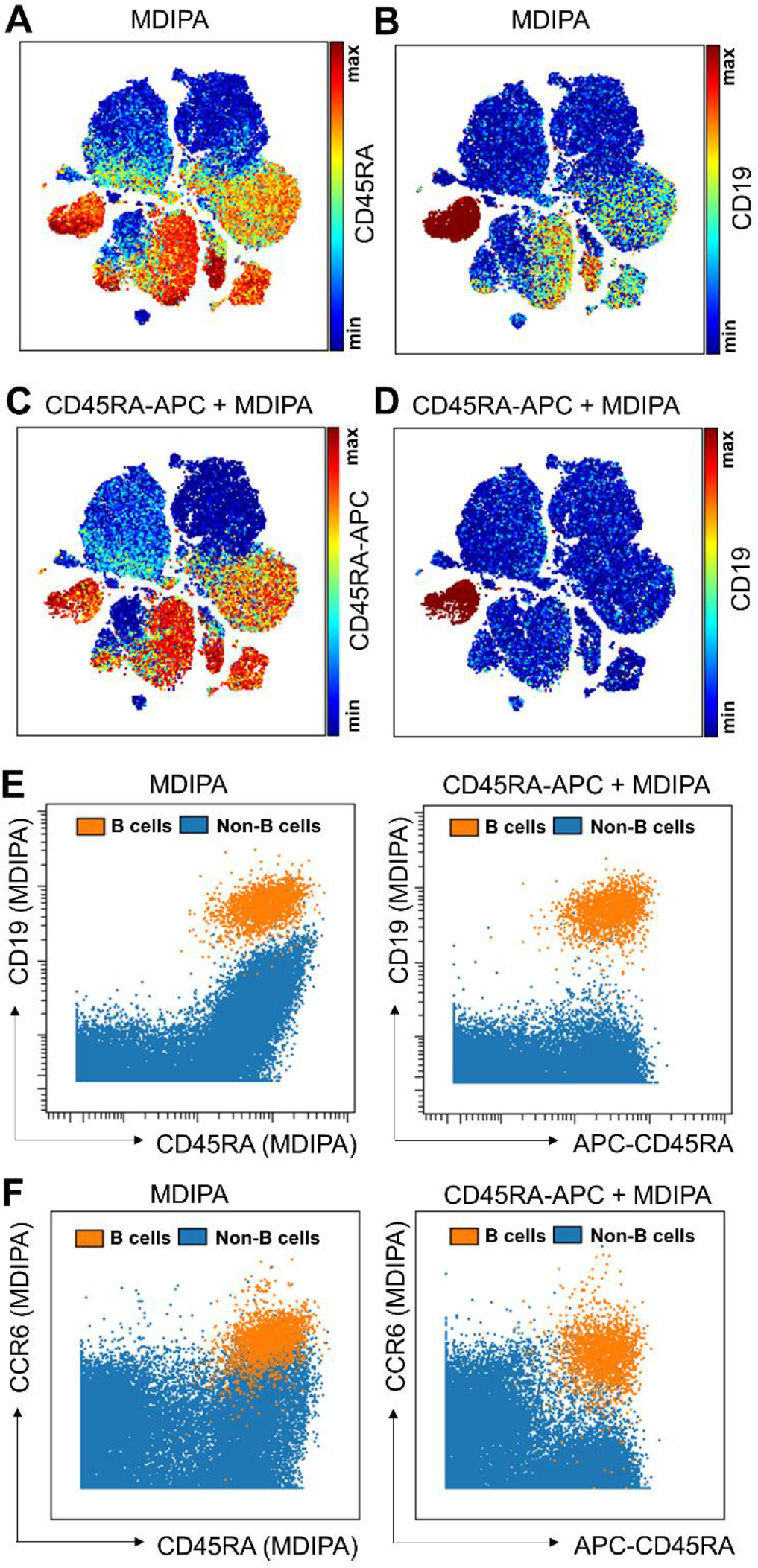
The MDIPA CD19 staining artifact is mediated by an interaction with CD45RA. (A-B) We observed that the aberrant CD19 staining was positively correlated with expression of CD45RA. (C-D) Whole blood from Donor A was co-stained with the MDIPA panel and a matching clone of APC-conjugated CD45RA to compete with the CD45RA antibody in the MDIPA panel. (C) Detection of the bound CD45RA-APC antibody with an anti-APC antibody shows that the competing antibody reproduces the original CD45RA expression pattern. (D) Competing out the CD45RA in the MDIPA panel also completely eliminates the aberrant CD19 staining. (E) Biaxial plots showing the correlated expression of aberrant CD19 expression with CD45RA in the MDIPA panel, but absence of correlated aberrant signal with the CD45RA-APC antibody. (F) Biaxial plots showing the correlated expression of aberrant CCR6 expression with CD45RA in the MDIPA panel, but absence of correlated aberrant signal with the CD45RA-APC antibody.

## Discussion

Flow and mass cytometry-based immunophenotyping relies on the assumption that antibodies bind specifically to their respective antigens, and that measuring the amount of bound antibody can therefore serve as an effective surrogate for measurement of the corresponding protein. In the case of non-specific antibody staining, this assumption no longer holds true, and antibody-associated signals can be erroneously interpreted and confound data analysis. There are several known sources of non-specific antibody signals in mass cytometry, which can be addressed in distinct ways: 1) antibody oxidation and isotopic impurities can result in signal cross-talk into adjacent or +16 mass channels, but this is a highly predictable phenomenon that can most effectively be addressed by optimizing panel design [7]; 2) non-specific staining can occur when certain cell types also bind antibodies through their Fc receptors, which can be mitigated by blocking Fc receptors prior to staining [8]; 3) non-specific staining can result from charge-based interactions of metal-conjugated antibodies with cationic granule proteins in eosinophils, which can be effectively blocked with heparin as a competing anionic protein [9]. Here, we report an entirely distinct form of non-specific antibody staining that appears to be more complicated, less predictable and which we have only observed when using the Fluidigm MDIPA panel to stain whole blood samples. We also note that while most of the data presented here were generated using a SmartTube-based adaptation of the MDIPA protocol, we have observed the same donor-dependent aberrant staining patterns when staining samples using the standard MDIPA protocol, and have also observed them in blood collected in sodium heparin and CPT citrate tubes (data not shown), suggesting that the blood collection tube and staining protocol are not major contributors to the issue.

Our data clearly demonstrate that this is a donor-dependent phenomenon that is specifically mediated by factors found in the plasma of specific donors. However, we did not find demographic associations between donor age, sex or ethnicity that might account for why some individuals are susceptible to the phenomenon and others are not. It is interesting to note that neither CD19 nor CD45RA have well defined physiological ligands, and both are heavily glycosylated proteins raising the possibility of interactions with lectins or other unknown ligands in the plasma. In an attempt to identify potential contributing factors, we evaluated data from a cohort of approximately 80 patients for whom we had performed matched CyTOF profiling of whole blood using the MDIPA panel and plasma proteomics using the SomaLogic SomaScan platform, which utilizes an aptamer based approach to evaluate the expression of thousands of plasma proteins [10]. However, we were unable to find any consistent correlations between the presence of the MDIPA CD19 staining artifact and any of the over 5000 proteins measured by the SomaScan assay (data not shown). Thus, the specific identity of the plasma factors that contribute to the aberrant staining phenomenon remain unclear.

It is also interesting to note that the aberrant staining patterns observed with the MDIPA are restricted to only specific antibodies in the MDIPA panel, namely, CD19, CCR6 and CD45RA. We noted that the staining intensity of CD19-144Nd in the MDIPA kit is much higher than the intensity of the same CD19 clone purchased as a pre-conjugated liquid antibody from Fluidigm or conjugated in house using X8 MaxPar polymer kits. While Fludigm has not disclosed the specific polymers used in the MDIPA kit, the higher signal intensity could suggest the use of novel higher-yield polymers, which could potentially be contributing to the phenomenon observed here. This also raises the potential that similar issues may potentially be seen outside of the MDIPA context when using liquid MaxPar antibodies that are conjugated to specific polymers, and we would recommend that Fluidigm disclose which polymers are used as part of the data sheets for their pre-conjugated antibodies to allow users to account for these potential issues.

While our studies have yet to define the exact mechanism underlying the aberrant antibody binding patterns, they do suggest several practical solutions that can help mitigate the problem. Given that the issue is caused by donor plasma, we do not anticipate any issues when applying the MDIPA panel to profile PBMC samples. If whole blood analysis is preferable for specific experimental reasons (e.g., to profile granulocytes), then depleting red blood cells and washing the leukocytes to remove residual plasma can effectively mitigate the problem. It should be noted that several of the antibodies in the MDIPA panel target fixation-sensitive antibody epitopes. Both of these approaches unfortunately add additional upstream processing steps that detract from the simplicity of the MDIPA whole blood assay. Given that the aberrant staining seems to be related to interactions between the CD45RA antibody in the MDIPA panel and the CD45RA antigen, another potential solution is to use the MDIPA together with a competing fluorophore-conjugated CD45RA antibody (e.g., CD45RA-FITC) to block binding of the CD45RA-150Nd in the MDIPA panel. After the blood has been washed and fixed, the fluorophore-conjugated CD45RA antibody can then be detected using a 150Nd-conjugated anti-fluorophore antibody, which will restore specific CD45RA staining back in the same channel it would originally have been present in while removing the artifactual staining. Finally, even if none of these steps can be taken, it is important for users of the MDIPA kit to be aware of this potential artifact and to screen samples for potential aberrant CD19 and CCR6 staining and exclude these parameters from analysis when necessary and to instead rely on other non-affected markers in the panel for cell identification (e.g., CD20 for B cells). Ultimately, we hope that Fluidigm is able to identify the specific cause of the aberrant staining patterns and to supply validated lots of the MDIPA kit to ensure reliable and reproducible whole blood CyTOF immunophenotyping.

